# The regulatory landscape of multiple brain regions in outbred heterogeneous stock rats

**DOI:** 10.1101/2022.04.07.487560

**Authors:** Daniel Munro, Tengfei Wang, Apurva S Chitre, Oksana Polesskaya, Nava Ehsan, Jianjun Gao, Alexander Gusev, Leah C Solberg Woods, Laura M Saba, Hao Chen, Abraham A Palmer, Pejman Mohammadi

## Abstract

Heterogeneous Stock (HS) rats are a genetically diverse outbred rat population that is widely used for studying genetics of behavioral and physiological traits. Mapping Quantitative Trait Loci (QTL) associated with transcriptional changes would help to identify mechanisms underlying these traits. We generated genotype and transcriptome data for five brain regions from 88 HS rats. We identified 21,392 cis-QTLs associated with expression and splicing changes across all five brain regions and validated their effects using allele specific expression data. We identified 80 cases where eQTLs were colocalized with genome-wide association study (GWAS) results from nine physiological traits. Comparing our dataset to human data from the Genotype-Tissue Expression (GTEx) project, we found that the HS rat data yields twice as many significant eQTLs as a similarly sized human dataset. We also identified a modest but highly significant correlation between genetic regulatory variation among orthologous genes. Surprisingly, we found less genetic variation in gene regulation in HS rats relative to humans, though we still found eQTLs for the orthologs of many human genes for which eQTLs had not been found. These data are available from the RatGTEx data portal (RatGTEx.org) and will enable new discoveries of the genetic influences of complex traits.

## Introduction

Rats are used in a variety of fields including physiological and behavioral research because of their similarities to humans and are preferred over mice for studying certain traits (1–4). In particular, research into the genetic basis of complex behavioral tasks such as measures of impulsivity and complex models of substance abuse and other motivated behavior has made extensive use of various inbred and outbred rat populations (5). Many associations with these traits have been detected (6). However, similar to the situation in human complex trait genetics, resolving implicated chromosomal regions to specific genes and underlying mechanisms is a critically important step that remains challenging.

Identification of heritable differences in gene expression via expression quantitative trait loci (**eQTL**) mapping offers one way to identify the molecular mediators of loci implicated by genome-wide association studies (**GWAS**) (7–10). Mapping of eQTLs has been conducted at scale for dozens of human tissues, most notably by the Genotype-Tissue Expression Consortium (**GTEx**) (7). In contrast, eQTL mapping in rats has been limited in terms of populations, tissues, sample size, and number of genetic markers used (11–27). Some eQTL mapping has been conducted in Heterogeneous Stock (**HS**) rats (28, 29), but to our knowledge this has not yet been done transcriptome-wide.

HS rats were developed in the 1980s by interbreeding eight inbred rat strains (ACI/N, BN/SsN, BUF/N, F344/N, M520/N, MR/N, WKY/N, and WN/N) (30) and have been maintained as an outbred population ever since. As a result, each HS rat chromosome is a mosaic of the eight possible founder haplotypes meaning that all alleles are common. The relatively high minor allele frequency is in stark contrast to humans, which have a preponderance of rare variants, and provides greater power for mapping eQTLs.

Because HS rats are being used for a variety of behavioral and physiological studies, there is an urgent need for a well-powered and complete library of QTLs. Here, we used tissue from five brain regions that have been implicated in addiction and other psychiatrically important traits to map eQTLs and splicing QTLs (**sQTLs**) in HS rats. We explored several important considerations for eQTL mapping in this population. We also have compared the results of eQTL mapping in HS rats to publicly available human data, identifying both similarities and important differences. Finally, we have provided all the data generated here on an online portal (RatGTEx.org) that provides a clearing house for this and other eQTL datasets.

## Materials & Methods

### Brain samples

Brains were extracted from 88 HS rats (43 male and 45 female). Mean age was 85.7 ± 2.2 for males and 87.0 ± 3.8 for females. Rats were selected to try to avoid related individuals. All rats were group housed under standard laboratory conditions and were naïve to behavioral or drug treatment.

Rat brains were taken out of a -80°C freezer and cryosectioned into 60 μm sections, which were mounted onto RNase-free glass slides. Slides were stored in -80°C until dissection. During dissection, slides were placed on a -20°C cold plate. One drop (approximately 50 μl) of RNAlater was placed on the brain region of interest. Each brain region then was dissected out under a dissecting video camera by using a pair of fine-tipped forceps with the assistance of an 18 gauge needle with a bent tip. Bilateral tissue of the same brain region from each rat was immediately transferred into 350 μl Buffer RLT (containing beta-mercaptoethanol) and placed on dry ice. Tissue was stored in -80°C before RNA extraction.

Tissue was thawed on ice and homogenized by using a clean stainless steel bead using Qiagen TissueLyser (40 Hz, 3 min). AllPrep DNA/RNA mini kit (Qiagen) was used to extract RNA. Samples were processed by using the QIAcube robot following standard protocols. The optional DNase digestion step was included for RNA samples. The average RIN for PL, IL, OFC, NAcc and LHb were 9.47 ± 0.58, 9.33 ± 0.63, 9.7 ± 0.53, 8.88 ± 0.79, and 8.94 ± 0.88, respectively.

### RNA sequencing

We performed RNA-Seq on mRNA from each brain region sample using Illumina HiSeq 4000 to obtain 100 bp single-end reads for 435 samples, with 26.7 million raw reads per sample on average (**Supplementary Table 1**).

To quantify gene expression, reads were first trimmed for adapter and poor-quality base calls using cutadapt (31). Reads were then aligned to the Ensembl Rat Transcriptome using RSEM (32). Upper quartile adjustment was applied to estimated gene read counts using DESeq2 (33). Samples were filtered based on low reads counts, mismatched genotypes (as described in the paragraph below), and expression principal component analysis (PCA) outliers. For two rats, all samples were removed by these filters, yielding processed data for 397 samples in 86 rats. Genes were eliminated if less than 25% of libraries had more than one read or if the total number of reads among all libraries for the gene was less than 100. Read counts were log_2_ transformed after adding a pseudocount of one to each read count. We used those values for calculating allelic fold change, and for eQTL mapping we applied rank-based inverse normal transformation to the values per gene.

Separately, to quantify allele specific expression and splicing, RNA-Seq reads were aligned to the Rnor_6.0 (rn6) genome from Ensembl (http://ftp.ensembl.org/pub/release-99/fasta/rattus_norvegicus/dna/Rattus_norvegicus.Rnor_6.0.dna.toplevel.fa.gz) using STAR v2.7.3a (34). STAR was run in two passes per sample, where novel splice junctions identified in the first pass were used to align additional reads in the second pass. The second pass used WASP to reduce mapping bias due to polymorphisms (35). Duplicate reads were then marked with the Picard MarkDuplicates function.

To check for mismatched RNA-Seq/genotype samples, we counted reads containing each allele for each exonic SNP using GATK ASEReadCounter (36) and compared counts to the genotypes at those SNPs. We identified 13 samples in which the RNA-Seq sample did not correspond to the label-associated genotype. Two of these samples matched each other’s genotypes, and their rat IDs were swapped and the samples were kept. Of the remaining 11, three matched with genotypes for which samples already existed for the same brain region, and the other eight matched with none of the 88 genotypes, so these 11 samples were removed from the study.

### Genotyping

We used genotyping-by-sequencing as described previously (37) to genotype the 88 rats, yielding 125,686 high-quality observed autosomal SNPs in Rnor_6.0 coordinates. We used SHAPEIT (38) followed by IMPUTE2 (39) to impute additional SNPs based on the genotypes of the eight HS founder strains (ACI/N, BN/SsN, BUF/N, F344/N, M520/N, MR/N, WKY/N, and WN/N), resulting in phased genotypes for 3,511,003 SNPs.

### Founder haplotypes

Regions of the 88 rat genomes were mapped to the eight HS founders using the calc_genoprob function of R/qtl2 with the cohort and founder strain genotypes (40). Diploid haplotype pair probabilities were collapsed to probabilities per strain per locus per animal using the genoprob_to_alleleprob function. These inferred haplotype mappings were used solely to examine genetic diversity in the cohort, and were not utilized in QTL mapping.

Haplotype probabilities were compared to the results of breeding simulations. For one simulated locus, one of eight haplotype labels was randomly chosen for each of the two copies per individual, for 100 individuals grouped into 50 female-male pairs. To progress one generation, individuals were rearranged into new pairs either by rotating the males by one in the sequence of pairs (circular mating), or by shuffling the pair assignment of the males (random mating). For each new pair, the locus is inherited in an offspring by randomly selecting one of the two alleles from the female and another from the male. This was done twice per pair to produce a new set of 50 female-male pairs. This was repeated for 80 generations. This full locus simulation was repeated 200 times using circular mating and 200 times using random mating.

### eQTL mapping

We performed cis-eQTL mapping using single-SNP linear regression implemented in tensorQTL (41), testing variants within 1 Mb upstream and downstream of each gene’s transcription start site (cis-window). We included 28 covariates: the first 20 principal components of the brain region’s expression matrix, and the genotype similarity to each of the eight HS founder strains to control for unequal relatedness. Empirical beta-approximated p-values were computed using data permutations (42) and were then used to calculate gene-level q-values and nominal p-value significance thresholds. A q-value cutoff of 0.05 was used to determine the genes for which at least one significant cis-eQTL was found. We then ran tensorQTL in cis_independent mode to find additional, conditionally independent cis-eQTLs per cis-eQTL gene (eGene) using a stepwise regression procedure (43). Finally, we ran tensorQTL in trans mode without excluding cis-window SNPs to identify all associations genome-wide with nominal p-value < 10^−5^.

### sQTL mapping

We quantified splice phenotypes by first identifying splice junctions using regtools (44). Using the cluster_prepare_fastqtl.py script provided by the GTEx pipeline, we clustered introns using LeafCutter (45), mapped clusters to genes, and applied filtering and normalization. We then mapped cis-sQTLs using tensorQTL, using genes as phenotype groups when doing permutations to compute empirical p-values. As with cis-eQTLs, a q-value cutoff of 0.05 computed across genes was used to determine significant cis-sQTLs, and used stepwise regression to find additional conditionally independent cis-sQTLs for each gene. We used the same eight genotype covariates as for eQTL mapping, plus the first ten principal components of the splice phenotypes.

### eQTL effect size

We define cis-eQTL effect size as allelic fold change (aFC) and computed it in two independent ways. Primarily, we computed aFC from total gene expression based on the additive cis-regulatory model (46), with the same covariates as were used for eQTL mapping. For validation, we computed haplotype-level allele specific expression (ASE) using phASER (47), which we then used to compute aFC for genes with sufficient ASE information (48). This method relies on phased genotypes and the SNPs detected in RNA-Seq reads.

Effect sizes for GTEx eQTLs were obtained from tables downloaded from https://gtexportal.org. Human-rat ortholog pairs were obtained from Ensembl BioMart.

### GEMMA

To assess the impact of using a linear mixed model (**LMM**) for mapping cis-eQTLs, the leave one chromosome out (LOCO) method was used, so GEMMA (49) was run in gk mode to create 20 kinship matrices, each based on all genotypes except those on the same chromosome as the genes for which it would be used. GEMMA was run on the nucleus accumbens core samples in lmm mode using the Wald test. It was run separately for each gene, testing only the gene’s cis-window variants with minimum minor allele frequency (MAF) = 5%. As with the tensorQTL mapping, the first 20 expression PCs were used as covariates, but the eight genotype-based covariates were omitted so as not to interfere with the random effect term. Percent variance explained (**PVE**) by the kinship matrix for each gene was computed by running GEMMA in vc mode, supplying a kinship matrix but not genotypes. The lmm mode mapping was repeated in lm mode for comparison, identically aside from not supplying a kinship matrix. These results were used only for the analysis on LMM impact, while results from tensorQTL described earlier were used for the remainder of the study.

### Heritability estimates

The cis-heritability (h^2^) for rat genes were calculated with GEMMA by first computing a kinship matrix for each gene using only cis-window variants. GEMMA was then run in vc mode, supplying the gene-specific kinship matrix and the same covariates used for eQTL mapping, and recording the PVE from the output log. Human cis-heritability estimates were previously computed (50).

### V^G^ estimates

V^G^ (the expected variance in the gene dosage due to interindividual genetic differences observed in allele specific expression) was estimated for each gene in each rat brain region by running ANEVA (51) using the phased, gene-level allele specific expression counts. V^G^ estimates for human genes were similarly obtained using GTEx v8 data, and V^G^ estimates calculated with at least 5000 ASE counts were included.

The human gene sets used to subset human-rat ortholog pairs for V^G^ comparison were based on those previously collected by Mohammadi et al. (51–53). We removed sets that overlapped with fewer than 20 ortholog pairs, and replaced the GWAS-derived sets with sets of all author-reported genes for traits in the GWAS Catalog v1.0.2 (54), choosing the 10 traits with the most author-reported genes while avoiding redundant traits.

### Variant annotation

SNPs were annotated with functional categories using the Ensembl Variant Effect Predictor (55). The background SNP set for enrichment was all cis-window SNPs for all tested genes. The test sets for enrichment were the cis-eQTL SNPs (eSNPs) with lowest p-value per eGene in each brain region, including multiple SNPs in the case of tied p-values.

### Colocalization

We collected linear association statistics from eQTL mapping in the five brain regions and from the nine traits from a published GWAS in HS rats (56). GWAS scores were available for a set of pruned SNPs (r^2^ < 0.95), so for each brain region we selected the top cis-eSNP per gene that was present in the pruned GWAS dataset to test for colocalization. We computed z-scores for eQTL and GWAS associations by dividing the slope by its standard error for each selected SNP. Using the summary data-based Mendelian randomization (SMR) method (57) we computed the approximate L^2^ test statistic and computed a p-value using the upper tail of the chi-squared distribution with one degree of freedom. We computed SMR p-values for each selected SNP and used a Bonferroni threshold to determine SNPs with significant colocalization. We repeated this for each of the 45 tissue-trait combinations.

## Results

We obtained gene expression profiles from five brain regions from 88 HS rats using RNA-Seq with an average library size of 26.7 million raw reads (**Supplementary Table 1**). The regions examined were: nucleus accumbens core (**NAcc**), infralimbic cortex (**IL**), prelimbic cortex (**PL**), orbitofrontal cortex (**OFC**), and lateral habenula (**LHb**) (**Fig. 1a**). These brain regions were selected because of their relevance to a variety of behavior traits, including but not limited to substance abuse-related traits.

**Figure 1:**
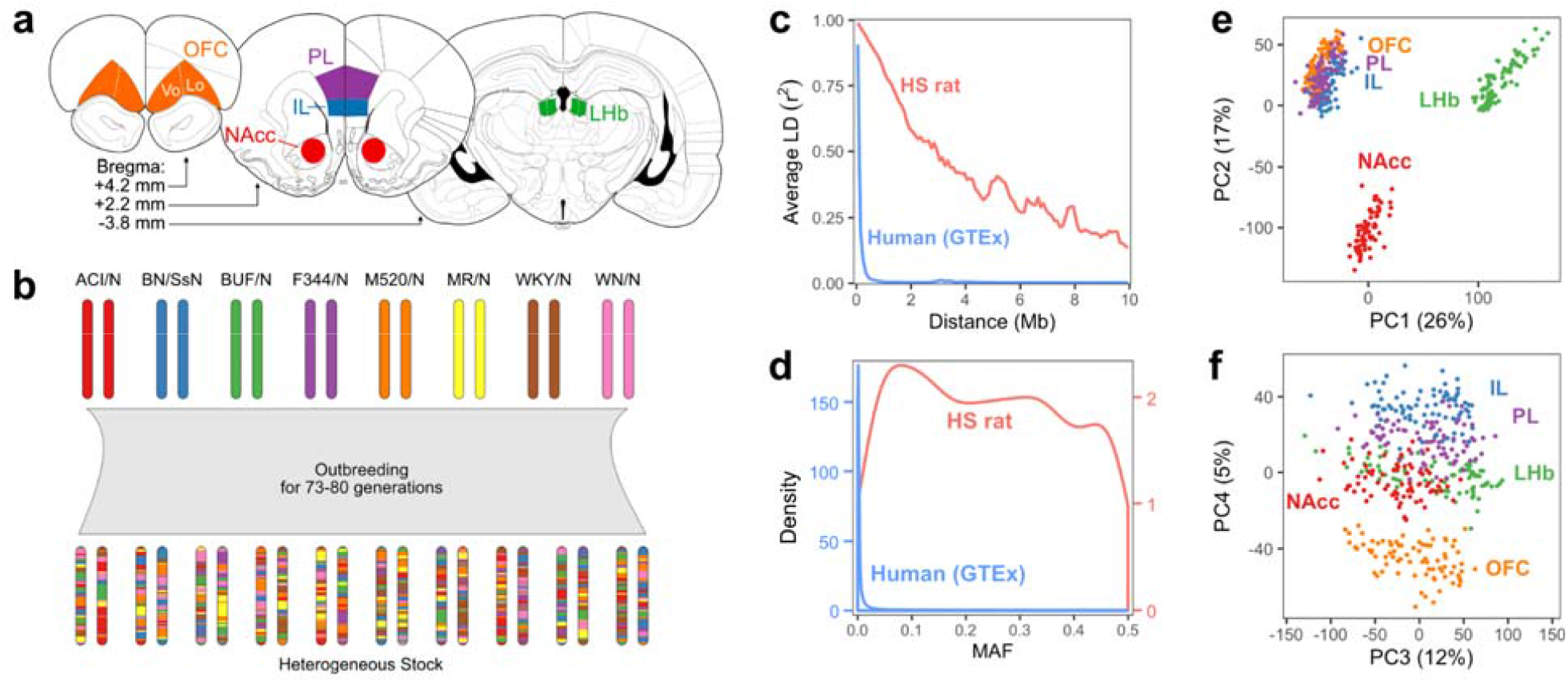
Overview of genotypes and expression data. **a** Diagram of the rat brain showing the five regions sampled in this study. **b** Diagram of the genetic outcome of heterogeneous stock rat breeding. The maternal and paternal chromosomes of a hypothetical autosome is depicted for each founder strain and for multiple HS rats. **c** LD decay in the HS rats in the present study and in humans in the GTEx project. LD was calculated using frequency-matched SNPs with MAF > 20%. **d** Distribution of SNP minor allele frequency in HS rats and in the humans in GTEx. **e**-**f** First two (e) and next two (f) components from principal component analysis applied to log-count expression data for all samples. IL, infralimbic cortex; LHb, lateral habenula; NAcc, nucleus accumbens core; OFC, orbitofrontal cortex; Vo, ventral orbital area; Lo, lateral orbital area; PL, prelimbic cortex; PC, principal component.

We determined genotypes at 3,511,003 SNPs across all autosomes using genotyping by sequencing (37). Consistent with our expectations based on their population history (**Fig. 1b**), linkage disequilibrium (**LD**) decayed over much longer distances in this population as compared to humans (**Fig. 1c**). Minor allele frequencies were fairly uniform, with a mean of 24% and the first and third quartiles of 11% and 36%; importantly the spike of rare alleles typically observed in human populations was not present (**Fig. 1d**).

Clusters of expression profiles for the three cortical regions were relatively close along their first two principal components, while nucleus accumbens core and lateral habenula profiles formed separate clusters (**Fig. 1e**). Further separation between the cortical regions was apparent in the fourth principal component (**Fig. 1f**).

### HS founder haplotype diversity

Since the HS rats have been maintained as an outbred population for many generations (73-80 for this cohort), the chromosomes are expected to be random mosaics of the eight founder haplotypes. In addition to the accumulation of recombinations, which improves mapping resolution, genetic drift inevitably erodes haplotype diversity (**Fig. 2**). We inferred ancestral haplotypes across each animal’s genome, which showed that at many loci the founder haplotypes had deviated substantially from their initially uniform proportions (**Fig. 2a, Supplementary Fig. 1**). To determine if the observed loss of haplotype diversity was consistent with genetic drift versus other possibilities such as genotyping errors, breeding errors, or inadvertent selection for fitness and fecundity, we simulated the breeding history. In particular, since the HS population has undergone periods of both circular and random pair mating, we simulated both strategies separately. We found that the distribution of observed haplotype diversity, as measured by Shannon entropy, lies between that of the two simulated strategies at generation 80 (**Fig. 2b**), suggesting that the changes in haplotype frequency are broadly consistent with random genetic drift.

**Figure 2:**
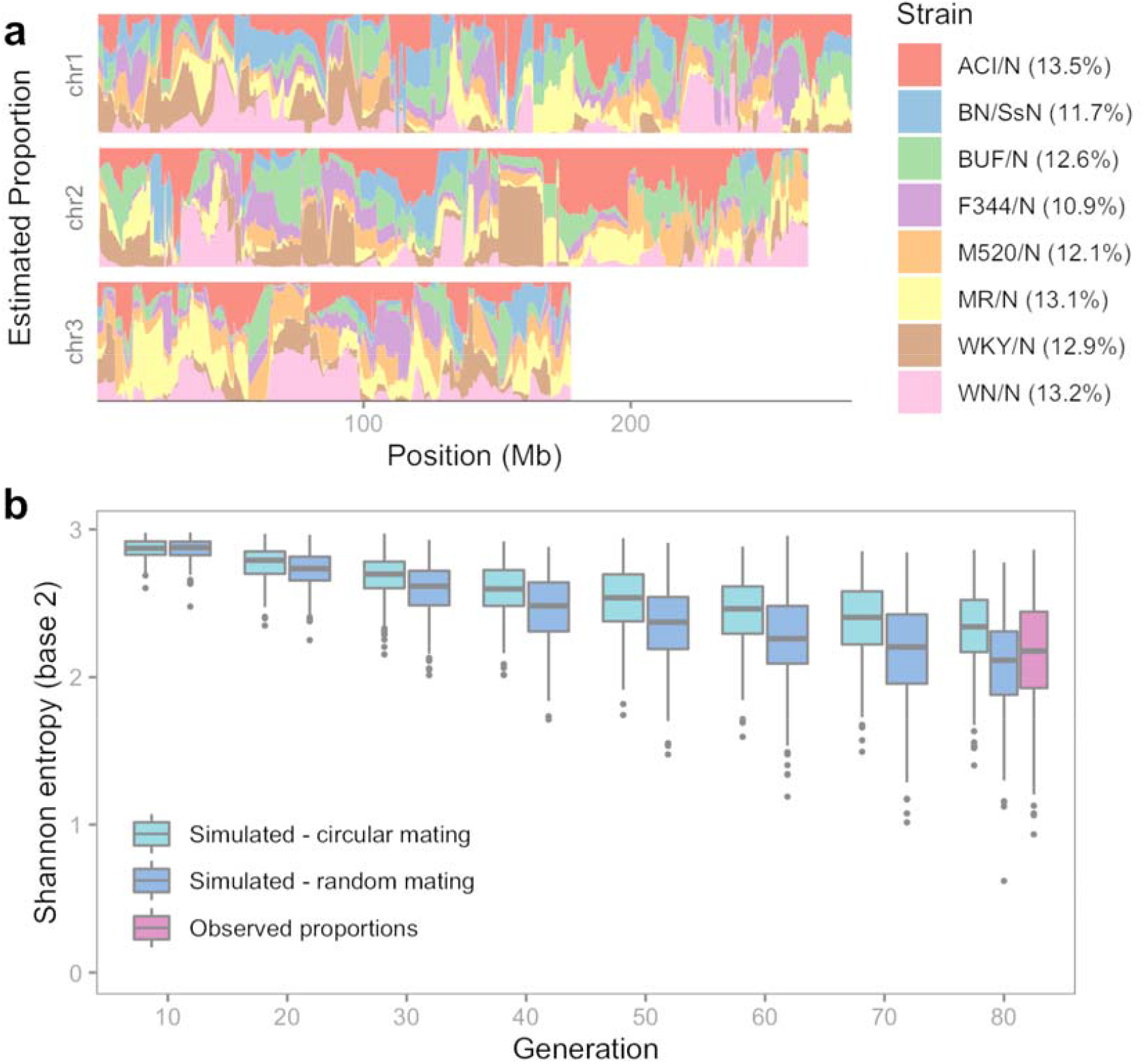
HS founder haplotype diversity. **a** Estimated haplotype proportions across three chromosomes in the 88 HS rats. The total proportions over all autosomes are shown in the legend. **b** Simulated and observed haplotype diversity in the HS population. Diversity (Shannon entropy of mean strain probabilities at each locus) is shown for every tenth generation for the simulated data. Each simulation type was run 200 times, representing 200 independent loci, and 200 real loci were sampled from the real cohort from HS generations 73-80.

### Mapping eQTLs

We tested for associations between gene expression and each SNP across the genome (**Fig. 3a**). As observed in other organisms (7, 9), associations with SNPs near the gene’s location in the genome, which we presumed to be cis-eQTLs, were prevalent. While we observed some associations with distant SNPs, which may represent trans-eQTLs, we primarily focused on putatively cis-acting eQTLs within ±1 Mb of each gene’s transcription start site (**TSS**) to retain statistical power and limit false positives (see Methods). Unless otherwise noted, “eQTL” hereafter refers to a cis-eQTL.

**Figure 3:**
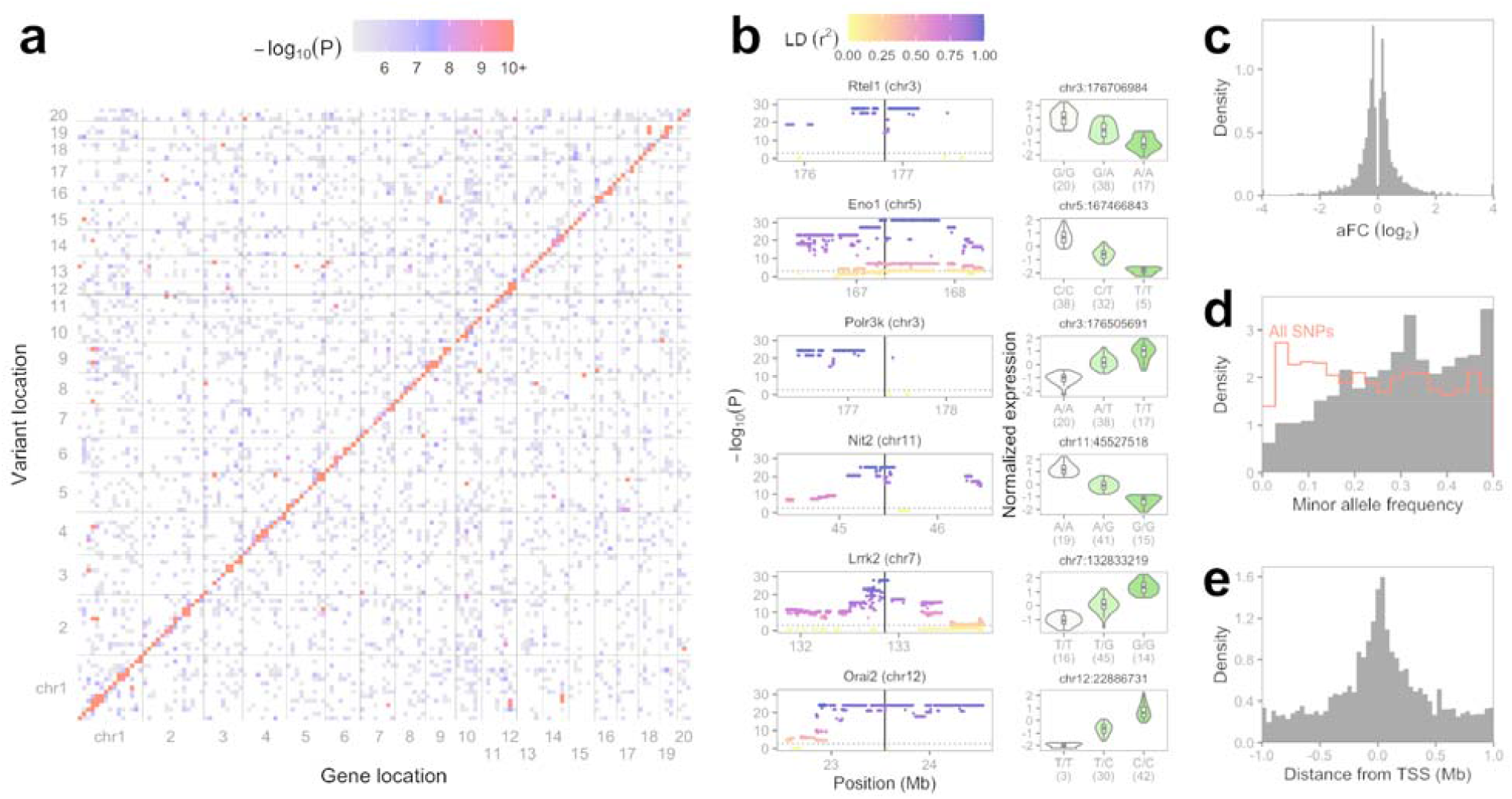
Overview of eQTLs for nucleus accumbens core. **a** Full-genome eQTL scans. Results are shown for nucleus accumbens core but were similar for all five brain regions. Each 20 Mb by 20 Mb bin is colored by its highest -log_10_(p-value), values below five are white. **b** Left: nominal -log_10_(p-values) for all cis-window SNPs for the top six NAcc cis-eQTLs, ranked by permutation-derived p-values. Vertical solid lines show each gene’s TSS, and horizontal dotted lines show each eQTL’s nominal p-value threshold as determined by permutations. The gene symbol and chromosome of each eQTL are labeled. LD values refer to each eQTL’s most significant eSNP(s). Right: effect plots for the representative eSNP per eQTL. Expression values are inverse normal transformed as used for eQTL mapping. Sample counts per genotype are shown in parentheses. **c** Allelic fold change of NAcc cis-eQTLs. **d** Minor allele frequency of top NAcc cis-eSNPs shown as solid gray bars. The red outline shows the histogram of minor allele frequency for all genotyped SNPs for comparison. **e** Top eSNP distance from TSS, oriented along the eGene strand. Each NAcc cis-eQTL is represented by one top eSNP, randomly chosen in the case of variants with identical p-values. TSS, transcription start site.

Plotting all p-values in the cis-windows revealed blocks of SNPs in full LD with identical p-values, or with multiple overlapping sets of such SNPs, depending on the founder haplotypes involved (**Fig. 3b**). The cis-window of 96.6% of the genes contained at least 10 SNPs that were not in full LD, and 28.1% contained at least 100 such SNPs. The TSS of 73% of genes were in high LD (r^2^ > 0.99) with at least one other gene’s TSS. In instances where multiple top SNPs in perfect LD were associated with the same gene, a single SNP was selected randomly for downstream analyses and visualization. We estimated the effect sizes for the cis-eQTLs using allelic fold change (**aFC**) and found that 87% of the cis-eQTLs alter the gene expression by up to two fold (|log_2_ aFC| ≤ 1), and 96% did so up to four fold (|log_2_ aFC| ≤ 2) (**Fig. 3c**). Consistent with their higher statistical power, minor allele frequencies of eQTLs were higher on average than the set of all measured SNPs (**Fig. 3d**). eQTLs were enriched close to the associated gene’s TSS, occurring upstream and downstream of the TSS at similar frequencies (**Fig. 3e**).

We identified cis-eQTLs for between 3,339 and 4,003 genes for each of the brain regions at a 5% false discovery rate (**Fig. 4a**), a consistent amount that represented 20% to 24% of the 16,456 to 16,814 expressed genes in each brain region. A total of 7,788 genes were affected by a cis-eQTL in at least one brain region, many of which (3,234, 42%) were identified in only one brain region and 1,170 (15%) of which were identified in all five brain regions (**Fig. 4b**). To validate the mapped cis-eQTLs, we measured their regulatory effect size from total gene expression and allele specific expression (**ASE**) data independently using aFC (46). We could obtain ASE counts for 70.8% of expressed genes on average per brain region, and from those counts estimated aFC from ASE for 52.4% cis-eQTLs on average. These two independent aFC measurements were consistent for each brain region (mean Pearson’s r = 0.58 ± SD 0.02, Deming regression β = 1.26 ± 0.10, **Supplementary Fig. 2**). We used a stepwise regression procedure to identify conditionally independent cis-eQTLs beyond the strongest cis-eQTL per gene, and found an average of 174 genes with two eQTLs and 4 genes with three eQTLs in each brain region, resulting in an average of 4.9% additional eQTLs per brain region. In total, we found 19,588 cis-eQTLs across the five brain regions (**Supplementary Table 2**).

**Figure 4:**
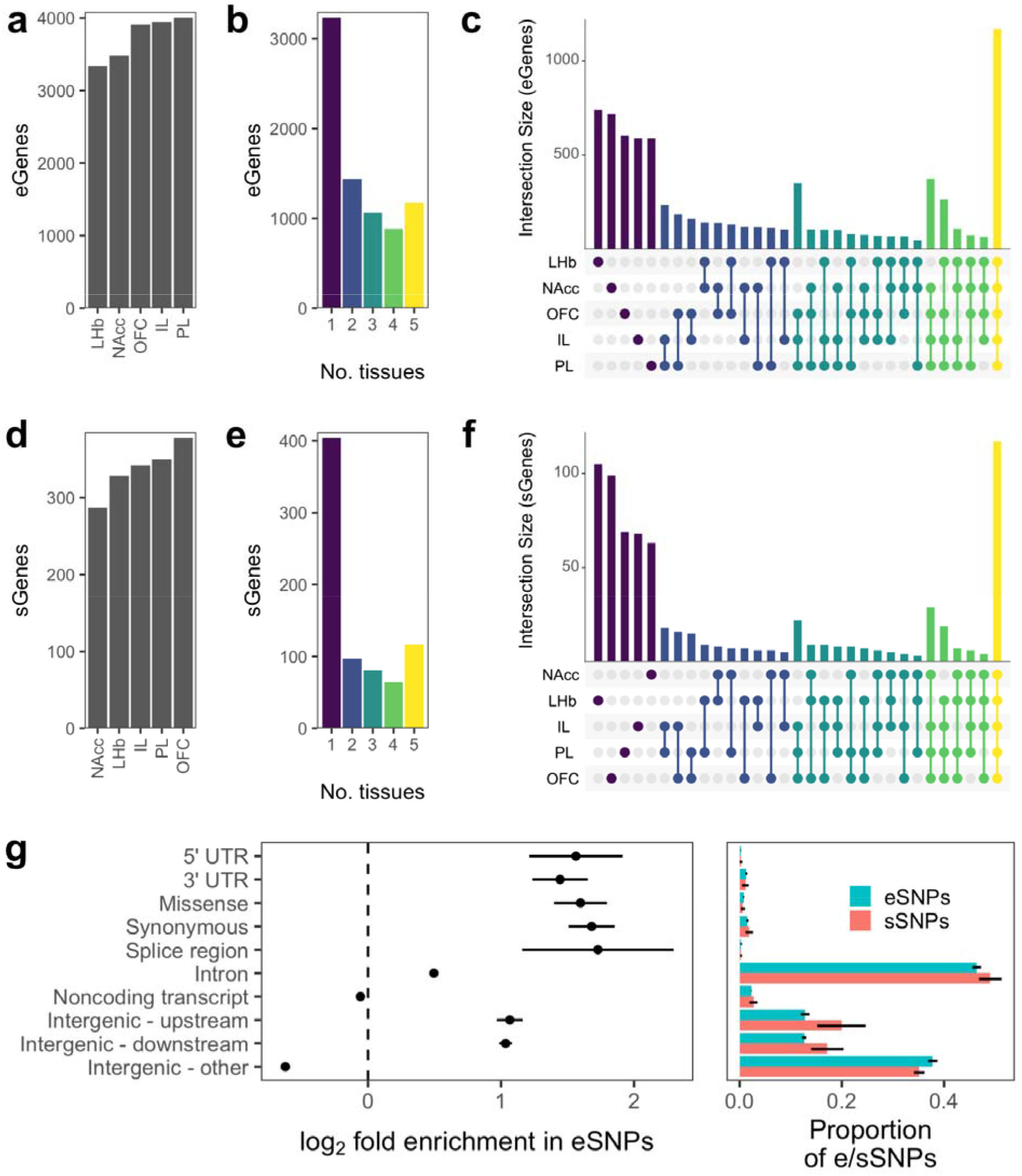
eQTLs and sQTLs across brain regions. **a** For each brain region, the number of genes for which at least one significant cis-eQTL was found. **b** The unique genes in panel (a), grouped by the number of tissues in which a cis-eQTL was found for the gene. **c** Overlap of eGene sets across tissues. Size of intersection is given for each combination of tissues, colored by the number of tissues in the combination. **d** For each tissue, the number of genes for which at least one significant cis-sQTL was found. **e** The unique genes in panel (d), grouped by the number of tissues in which a cis-sQTL was found for the gene. **f** Overlap of sQTL gene (sGene) sets across tissues. **g** Left: Enrichment of functional annotations in eSNPs (top associated SNP per cis-eQTL). Right: Proportion of eSNPs and sSNPs (top associated SNP per cis-sQTL) with each annotation. Representative eSNPs and sSNPs were randomly chosen among the top associations per e/sQTL in case of ties. Some SNPs have more than one annotation. Enrichment is with respect to all tested (cis-window) SNPs. Points and bars show mean across the five brain regions, and lines show standard deviations.

Next we looked at the tissue specificity of the eQTLs that may reflect biological differences across brain regions. The three cortical regions (IL, PL, and OFC) shared a greater overlap of cis-eQTL genes (**eGenes**) than any other tissue trio, with 349 eGenes shared exclusively among them, compared to only 104 eGenes for the next-highest trio (**Fig. 4c**). This observation is broadly consistent with our expectation that cortical tissues should have similar expression patterns. However, there were a number of eQTLs that were only detected in a single tissue (**Supplementary Table 3**). In some cases, this may reflect real biological differences that give rise to tissue specific eQTLs. In other cases, this apparent tissue specificity could be also caused by noise in associations that are close to the significance threshold, such that only one tissue reached the significance threshold, or by low expression of the gene in question in other tissues (**Supplementary Table 4**).

We quantified splicing in terms of intron excision ratios and used these phenotypes to map cis-sQTLs (7). Because these measurements are based on a smaller number of reads, power to detect sQTLs is likely lower than for eQTLs. We found cis-sQTLs in 4.1% to 5.4% of the 6,918 to 7,676 genes in which we detected alternative splicing per brain region. Over all splice junctions per gene and using stepwise regression, we found 305 to 403 independent cis-sQTLs per brain region, impacting a total of 764 genes (**Fig. 4d, Supplementary Table 5**). This included 404 genes for which cis-sQTLs were identified in only one brain region and 117 for which cis-sQTLs were found in all five brain regions (**Fig. 4e, Fig. 4f**). Importantly, 47% of the genes with an sQTL in a brain region did not have an eQTL in that brain region, demonstrating the added benefit of mapping sQTLs.

We annotated top associated SNPs (**eSNPs**) per eQTL and found enrichment in all protein-coding gene-associated categories, both exonic and intronic (**Fig. 4g**). However, these enrichments may also reflect the tendency for both eSNPs and gene-associated features to occur near the gene’s transcription start site relative to the full cis-window. While there were too few sQTLs to reliably measure sQTL SNP (**sSNP**) annotation enrichment, especially for smaller categories such as “Splice region”, the proportions of annotations among the sSNPs were similar to the proportions among eSNPs. Given that there were often blocks of multiple eSNPs with identical p-values, the level of eSNP resolution in HS rats is limited by LD structure.

Because of the complex familial relationships among members of an outbred population like the HS, several prior eQTL mapping studies in similar mouse populations employed a linear mixed model (**LMM**) that includes a kinship matrix that accounts for relatedness (9, 58, 59). However, LMMs are more computationally intensive, and given the large number of genes being examined, we questioned the need for an LMM. We examined genetic relatedness between the rats and found several outliers that appear to be closely related pairs (**Supplementary Fig. 3a**). We repeated cis-eQTL mapping for one brain region, NAcc, using GEMMA (49) in linear model (**LM**) mode and in LMM mode, run with identical parameters aside from the inclusion of a leave-one-chromosome-out kinship matrix for LMM. The absolute values of Z-scores for the top association per gene were highly correlated (Spearman’s rho = 0.991, **Supplementary Fig. 3b**). The sets of eGenes below a wide range of p-value thresholds had strong overlap between the modes (97%, 97%, and 95% for 10^−9^, 10^−6^, and 10^−3^, respectively, **Supplementary Fig. 3c**). This is in stark contrast to the scenario of GWAS in a panel of inbred mouse or rat strains, where use of LMM is critical to avoid false positive results (60). In our dataset, p-values for LMM tended to be slightly more significant (**Supplementary Fig. 3d**).

### Comparison to human

The GTEx Consortium recently released a comprehensive map of eQTLs in 49 human tissues including 13 brain regions (7). The number of eGenes in our data was lower (mean of 3736 over five tissues) than for the human brain tissues in GTEx data (mean of 6870 over 13 tissues) where the authors used 114 to 209 donor samples to map eQTLs, using a similar testing procedure and the same false discovery rate (5%) as the present study. We sought to compare these counts in light of the correlation between sample size and eGene count among the GTEx tissues (Pearson’s r = 0.86, **Fig. 5a**). We subsampled each GTEx brain tissue dataset to 81 samples, the largest sample size in the present study. Mapping eQTLs with these subsampled datasets resulted in fewer eGenes (mean: 1900, SD: 473) than the rat brain tissues (mean: 3736, SD: 304). While this comparison pertains to human datasets with artificially reduced sample sizes, and the human and HS rat datasets differed in multiple biological and technical ways that could influence statistical power, it suggests that on a per subject basis we had greater power to map eQTLs in HS rats.

**Figure 5:**
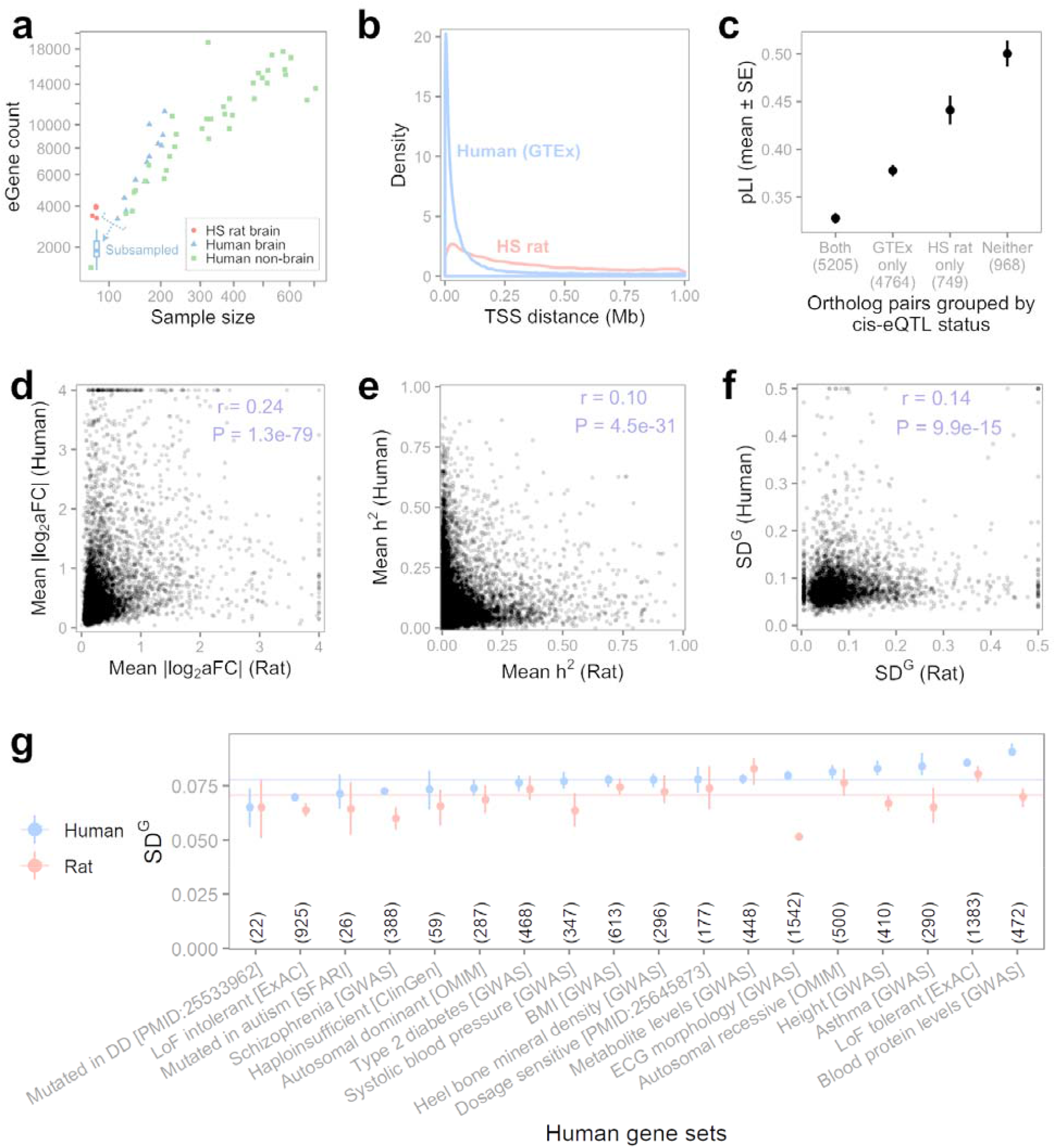
Comparisons to human eQTLs. **a** Scatter plot showing the relationship between sample size and number of detected eGenes for the five rat brain tissues in the present study and for all GTEx tissues. The boxplot summarizes the 13 counts for the human brain tissues after subsampling each dataset to 81 samples. Both axes are square-root-scaled. **b** Distributions of distances to transcription start site for each top eSNP in HS rat and GTEx brain tissues. **c** Probability of loss-of-function intolerance (pLI) (61) for expressed ortholog pairs in which one, both, or neither gene had an eQTL in any brain tissue. Ortholog pairs counts are shown in parentheses. **d** Pearson correlation of eQTL effect size in ortholog pairs in which eQTLs were found for both genes. For each gene, |log_2_(aFC)| for significant eQTLs were averaged across rat and across human brain tissues. **e** Pearson correlation of cis-heritability estimates in ortholog pairs. For each gene, h^2^ was averaged across all rat and all human brain tissues where it could be computed. **f** Pearson Correlation of SD^G^ 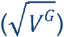 estimates in ortholog pairs. For each gene, SD^G^ was averaged across all rat and all human brain tissues where it could be computed. **g** For each set in a collection of human gene sets, averaged SD^G^ values from (f) subsetted to gene ortholog pairs in which the human gene is in the set. Points are medians, vertical segments are 95% confidence intervals of the medians, and horizontal lines show medians of all human or rat values from (f). Sets are sorted by human gene median, and set sizes, considering only the genes with SD^G^ data, are indicated in parentheses. DD, developmental disorder; LoF, loss-of-function; BMI, body mass index; ECG, electrocardiogram.

The distances between each eGene’s top eSNP and TSS were much greater for rat brain (median 271 Kb) than for human brain (median 35 Kb, **Fig. 5b**). In many cases a rat brain eQTL had a cluster of eSNPs in perfect LD (r^2^=1), which therefore had identical p-values. In these cases a single SNP was randomly chosen, which is one reason for the greater distances between the top eSNP and the TSS observed in HS rats.

Colocalization analysis and other transcriptome-informed functional population genomic analyses rely on presence of common regulatory variation in a population such as eQTLs to interpret GWAS signal. Next, we focused on genes that do not have an eQTL mapped in any brain tissues in the GTEx data. Out of 11,686 orthologous genes that are well expressed (median TPM > 1 in at least one tissue) in both GTEx human and HS rat data, we found that 85% have an eQTL in at least one GTEx brain tissue, leaving 1,717 genes with no mapped eQTLs. As previously reported, the genes with no eQTLs are enriched for critical genes that are intolerant to loss of function coding genetic variation (46). We found that for 44% (n=749) of these genes we identified an eQTL in at least one rat brain region. Indeed, the orthologous genes associated with these eGenes that are exclusive to the rat eQTL data are significantly more likely to be intolerant of loss-of-function mutations than those genes with eQTLs in GTEx data (**Fig. 5c**). These results suggest that the HS rat population may be a valuable resource for characterizing phenotypic consequences of genetic variation in genes that are highly depleted for functional variation in human populations.

### Conservation of genetic regulatory constraint between human and rat

Genetic regulatory variation present in a population is negatively correlated with the coding constraint of the genes (46, 51, 62). We compared the amount of genetic variation present in the HS rat population to human data using different approaches, each affected by a different set of confounding factors. Effect sizes (|log_2_ aFC|) of the cis-eQTLs in rat brain regions were smaller overall than those measured in 13 human brain tissues in GTEx (7). We looked at eQTL effect sizes for ortholog pairs to detect correspondence in tolerance to regulatory variation between similar human and rat genes. For each gene, we averaged the absolute effect size per top eQTL across tissues, and then paired up these values for every ortholog pair with any eQTL in rat brain tissues and any eQTL in human brain tissues (n = 6079 pairs). Effect sizes correlated significantly (Pearson’s r = 0.24, P = 1.3e-79, **Fig. 5d**). This suggests that some degree of variance in tolerance to regulatory variation is conserved between rat and human.

For each gene in each tissue, we estimated cis-heritability (**h**^**2**^) of expression. Since h^2^ is another measure pertaining to genetic regulatory constraint, we expected some correlation in h^2^ between orthologous genes due to their similarity in function and therefore correlation in their degree of evolutionary constraint. We averaged h^2^ across tissues per gene and compared to h^2^ estimates for human genes averaged over 13 GTEx brain tissues. Mean h^2^ between orthologs correlated modestly but was highly significant due to the large number of observations (Pearson’s r = 0.096, P = 4.5e-31, **Fig. 5e**).

We then estimated V^G^, the expected variance in the gene dosage due to interindividual genetic differences observed in allele specific expression using ANEVA (51). We compared the SD^G^ (standard deviation, 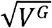) values per gene to those estimated for GTEx brain tissues. As expected, V^G^ correlated much more highly between tissues from the same species than did V^G^ between orthologs in cross-species tissue pairs (**Supplementary Fig. 4**). When averaged across tissues per rat gene and human gene, SD^G^ values for ortholog pairs were weakly but significantly correlated (Pearson’s r = 0.14, P = 9.9e-15, **Fig. 5f**). SD^G^ tended to be lower (i.e., lower genetic dosage variance) for the rat gene in ortholog pairs, including the orthologs for a wide range of human gene sets representing both essential and non-essential genes (**Fig. 5g**).

### Colocalization

Due to the longer-range LD in HS rats compared to humans, particularly the blocks of SNPs in complete LD within the cohort, colocalization methods that model colocalization as the overlap of single causal SNPs are less informative because colocalization probabilities are divided among the group of SNPs that are in LD with one another. To address this limitation, we tested colocalization of cis-eQTLs with GWAS results for a set of nine traits related to body morphology and adiposity obtained from an independent cohort of HS rats (56) using the summary data-based Mendelian randomization (SMR) method, which only evaluates consistency of effect for the top eQTL (57). We found 80 significant colocalizations among the 45 tissue-trait pairs using tissue-trait-specific Bonferroni p-value thresholds ranging from 1.3e-5 to 1.6e-5 (**Supplementary Table 6**), with the most colocalizations found for prelimbic cortex eQTLs and the RetroFat trait (**Fig. 6a**). Eight eGenes were involved in at least four colocalizations: *Apip, Cacul1, Drc1, Gpn1, Mrpl45, Nudt4, Pnpo*, and *Rbks*. Colocalizations for multiple tissues and traits generally clustered together in or near the QTL regions of the original GWAS (**Fig. 6b, Supplementary Fig. 5**).

**Figure 6:**
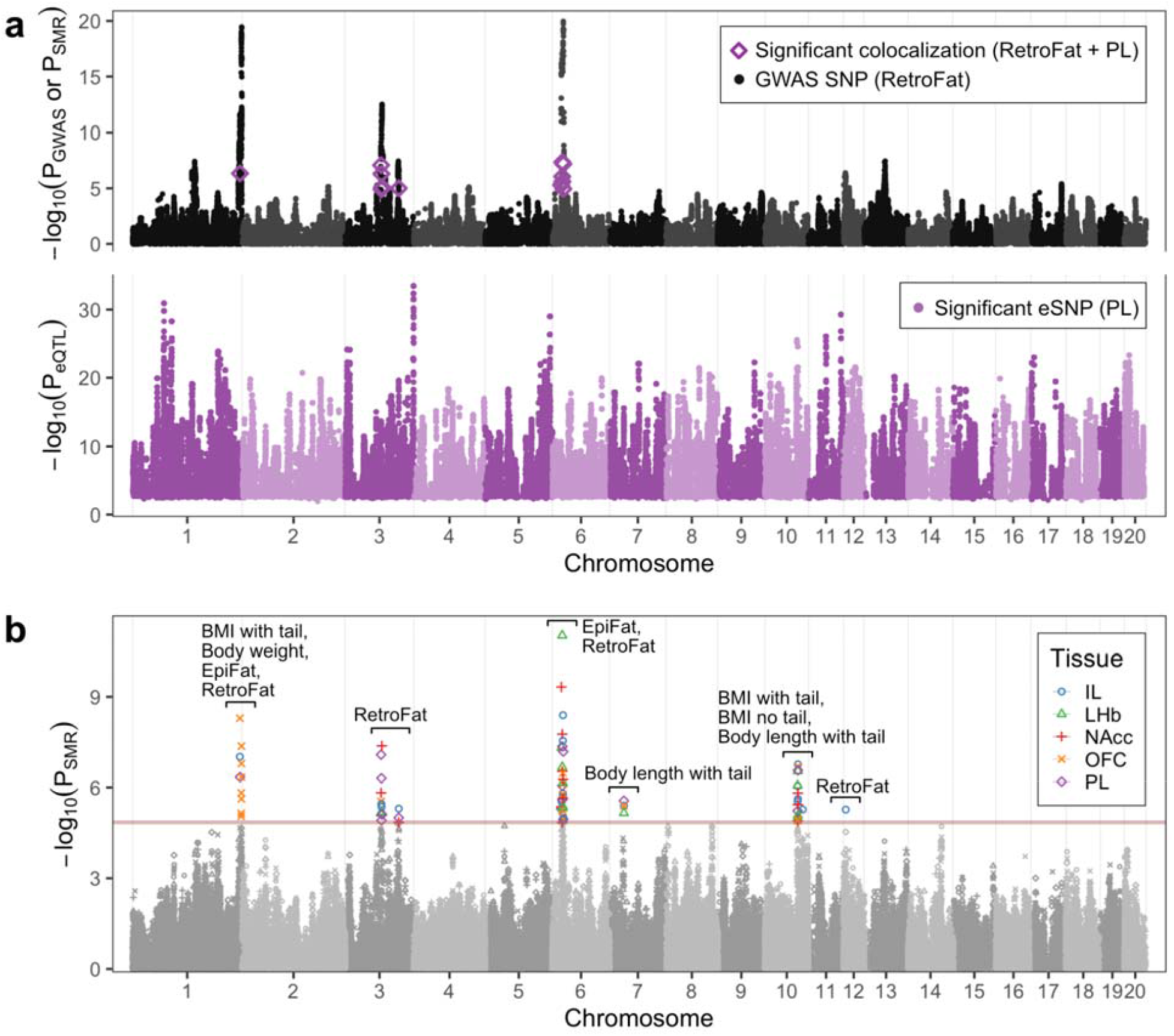
Colocalization of eQTLs and GWAS. **a** Colocalization for one tissue-trait pair, prelimbic cortex and RetroFat. Top, p-values from the RetroFat GWAS, with p-values for significant SMR tests overlaid. Bottom, p-values for significant eSNPs in PL. **b** SMR p-values for all tissue-trait pairs overlaid. Only significant SNPs are colored by tissue, and the remainder are gray. The traits involved in each cluster of significant colocalizations are labeled. Bonferroni p-value thresholds for each tissue-trait pair are shown as horizontal lines colored by tissue.

### Data portal

All gene expression, eQTL, and sQTL data are available at RatGTEx.org, for which we have adapted code and API design from the GTEx Portal to host rat eQTL data. This portal also includes interactive visualizations, derived from those in the GTEx portal, that can display results for any queried genes and variants. These five datasets initiate the RatGTEx portal, with datasets for additional tissues to be added as they become available.

## Discussion

We used RNA-Seq to map eQTLs and sQTLs in five brain regions in a cohort of 88 outbred HS rats. We also explored the unique genetic characteristics of the HS rat population. We focused on cis-eQTLs and sQTLs and characterized the degree of tissue specificity. We compared our results to human eQTL data from the GTEx project. We also used colocalization to demonstrate the utility of these eQTLs for interpreting GWAS results from HS rats. We have made all of the data generated here including the eQTL and sQTL mapping results available through a new portal (RatGTEx.org) to facilitate the application of these data to rat genetic and genomic research.

We mapped both cis-eQTLs and trans-eQTLs. The trans-eQTLs were much less prevalent. The biological significance of trans-eQTL signals is generally harder to ascertain as the analysis suffers from limited statistical power and can be confounded by batch effects. Furthermore, the rat genome assembly is not as thoroughly characterized as the human genome. Thus, some trans-eQTLs may reflect mismapping of reads from RNA-Seq such that a cis-eQTL appears to be a trans-eQTL or the information about a SNP’s location can also be incorrect due to an error in the genome assembly, which also creates an apparent trans-eQTL that is actually a cis-eQTL, as has been observed in human eQTL studies (63). For all of these reasons, we focused most of our efforts on cis-eQTLs. Future work with large sample sizes and a focus on trans-eQTLs could yield interesting results, for example, pertaining to colocalization analysis to understand pleiotropic GWAS QTLs.

We compared HS rat data presented here with human data from the GTEx project. We found fewer cis-eQTLs in rats compared to humans. This difference is consistent with our smaller sample size. Indeed, when we downsampled GTEx brain datasets so that the number of individuals matched our study, we only identified half as many eQTLs compared to our rat data. One reason that HS rats had more power than humans on a per-sample basis might be the longer-range LD in HS rats, which reduces the effective number of tests being performed (64). Another advantage of HS rats is the higher MAF as compared to humans. The greater power in HS rats could also reflect the much more controlled environment of laboratory rats.

The greater LD in HS rats compared to humans increases power but does so at the expense of precision since there are often large LD blocks that increase uncertainty about which SNP causes a given eQTL. Another consequence of this causal SNP uncertainty is that the eSNP annotation enrichments reported here are less indicative of the specific regulatory mechanisms driving the effect compared to those obtained in humans. For example, the distances between the eSNP and the TSS in rats is much wider in HS rats as compared to humans (**Fig. 5b**).

We found ten times fewer cis-sQTLs compared to cis-eQTLs. The GTEx project reported about four fold fewer cis-sQTLs compared to cis-eQTLs. The larger difference between sQTLs and eQTLs in our dataset may be due to both our use of single-end sequencing and our lower sequencing depth, both of which reduce the number of junction-spanning reads, which are essential for sQTL detection. Therefore, we do not believe this difference reflects a true biological difference between the two species.

Our study is similar to several previous studies that have mapped eQTLs or similar features in mice and rats. Prior mouse and rat studies have used inbred, recombinant inbred, and outbred populations, with microarrays or RNA-Seq (9, 11–29, 58, 65–70). While some of these previous studies have used more computationally intensive linear mixed models to account for population structure effects, we did not find appreciable difference between the results from the linear regression and linear mixed model, which is consistent with Parker et al. (9). This may be explained by the fact that we avoided sampling multiple individuals from the same family. Had the breeding scheme been less carefully designed, there may have been isolated clusters of more closely related individuals within the population, and an LMM might have been necessary.

Genes that are intolerant to loss of function mutations tend to have lower levels of regulatory variation as well (46). We found cis-eQTLs in the rat orthologs of many of the human genes with no cis-eQTLs in the GTEx brain tissue data. Indeed, these genes with eQTLs in only rats had relatively high intolerance scores in humans. Given the lower statistical power in our study versus the GTEx brain dataset, these rat exclusive eQTLs are likely a result of relaxed selection pressure against eQTLs in these genes in rats. Presence of common regulatory variants in these genes presents an opportunity to study the downstream dosage effects in some of these variation-intolerant human gene orthologs.

Comparing the amount of genetic variation in gene expression in rats and humans, we found that this quantity is only moderately correlated between the populations. This low correlation level is likely a combined effect of low statistical power and the artificial nature of the rat population that relaxes selection constraints on genes. However, surprisingly, we found that the rat population shows lower levels of genetic regulatory variation across a diverse set of genes as measured by eQTL effect sizes, cis heritability of gene expression, and the ASE-derived estimates of genetic variance in gene expression. Notably, these results cannot be explained by the difference in statistical power and the sample sizes. Future investigation could uncover the cause, in particular whether it relates to biological differences between humans and rats, consequences of the HS rat population design, environmental conditions, or other factors.

We were able to use eQTLs from brain tissue to show colocalization with nine body morphology and adiposity traits. The success of this approach may reflect the idea that eQTLs are shared across many tissues, not just among brain regions. Furthermore, adiposity is heavily influenced by consummatory behavior and energy expenditure, both of which are controlled by the brain.

The results of this study offer practical guidance for future HS rat eQTL studies. For example, the degree of eQTL overlap across brain regions was very high, especially for the three cortical regions. Had we sampled the fewer brain regions from a larger number of individuals, we would have obtained greater statistical power.

## Data availability

Raw RNA-Seq data is available at NCBI GEO accession GSE173141. Processed genotype, expression, eQTL, and sQTL data are available at https://RatGTEx.org/download/#studies.

## Acknowledgements

This work was supported by the National Institute on Drug Abuse (P50DA037844 and P30DA044223), the National Institute of General Medical Sciences (R01GM140287), and the National Institute on Alcohol Abuse and Alcoholism (R24AA013162). DM, PM, and LMS were partly supported by Skaggs Scholars Program.

## Author contributions

AAP and HC designed the study. LSW bred and shipped the animals. TW bred and processed the animals and produced RNA-Seq data. HC supervised tissue collection and conducted a preliminary analysis. JG helped to produce the genotypes used in our analysis. LMS processed RNA-Seq data and quantified gene expression. DM, ASC, OP, and NE processed RNA samples and analyzed data. PM, AAP, and AG designed and advised on data analyses. PM, AAP, and DM wrote the paper.

