## Supplemental Materials for "The regulatory landscape of multiple brain regions in outbred heterogeneous stock rats"

##### Supplementary Tables (.xlsx file)

**Supplementary Table 1.** Statistics for RNA-seq and quality control.
**Supplementary Table 2.** The cis-eQTLs identified in each brain region.
**Supplementary Table 3.** The genes with the strongest tissue-specific cis-eQTLs across the five brain regions.
**Supplementary Table 4.** The genes with the strongest tissue-specific expression across the five brain regions.
**Supplementary Table 5.** The cis-sQTLs identified in each brain region.
**Supplementary Table 6.** The significant colocalizations between cis-eQTLs and adiposity trait GWAS loci.

Supplementary Figure 1

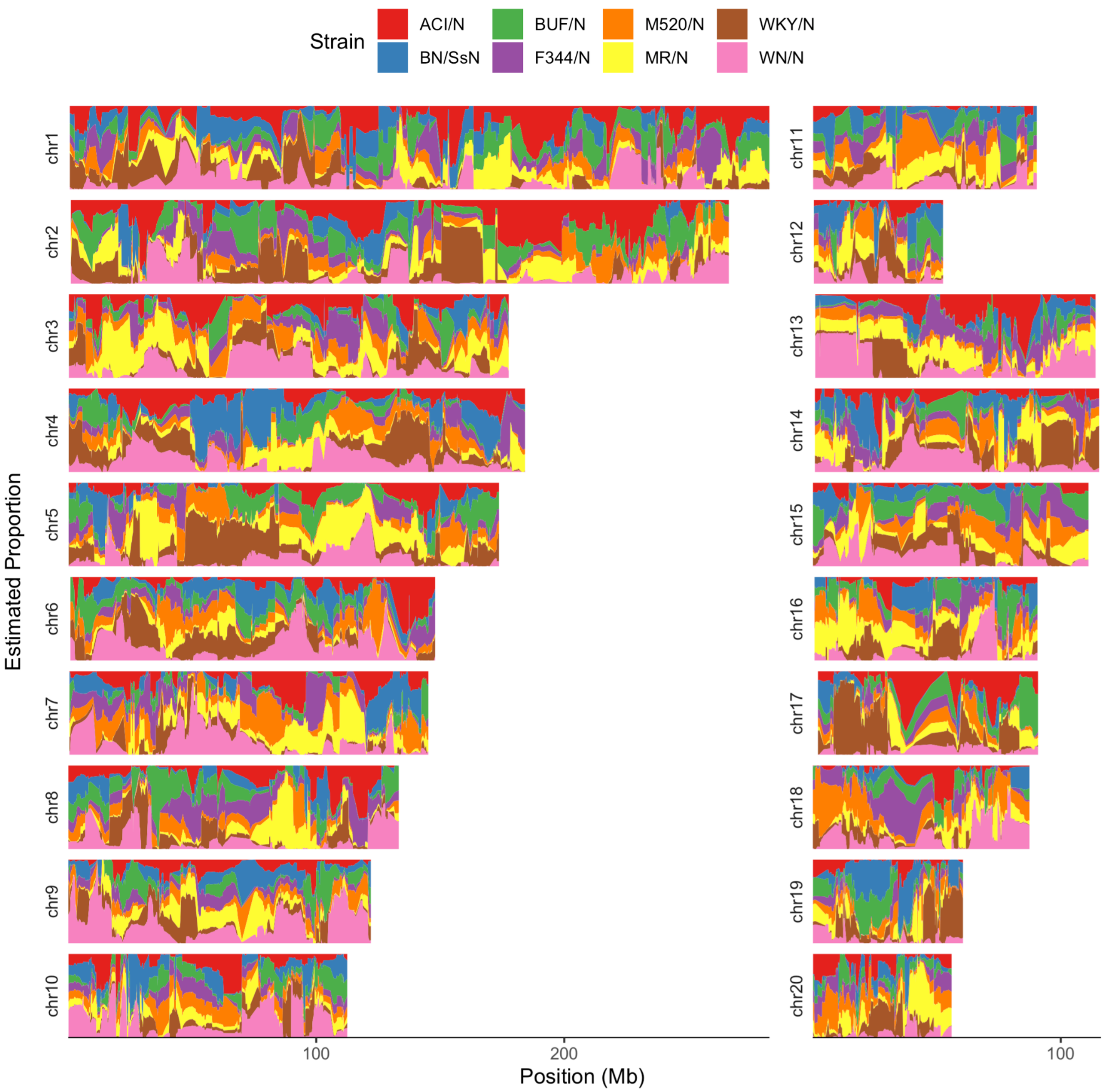


**Founder haplotype estimates.** Probability of descent per founder strain at each chromosome position, averaged over the 88 rats and the two haplotypes per rat.

### Supplementary Figure 2

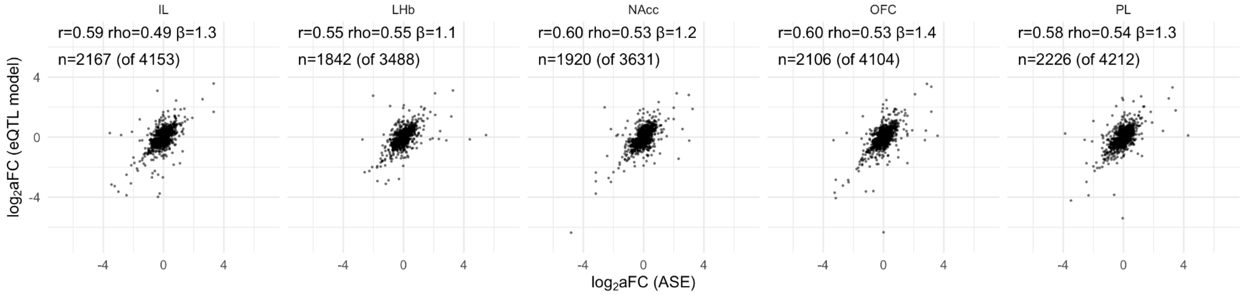


**Effect size agreement between eQTL model and allele specific expression.** Points are shown for the subsets of conditionally independent cis-eQTLs for which ASE could be measured, with n giving the size of these subsets out of the total cis-eQTL counts in parentheses. Coefficients for Pearson correlation (r), Spearman correlation (rho), and Deming regression (β) are shown.

Supplementary Figure 3

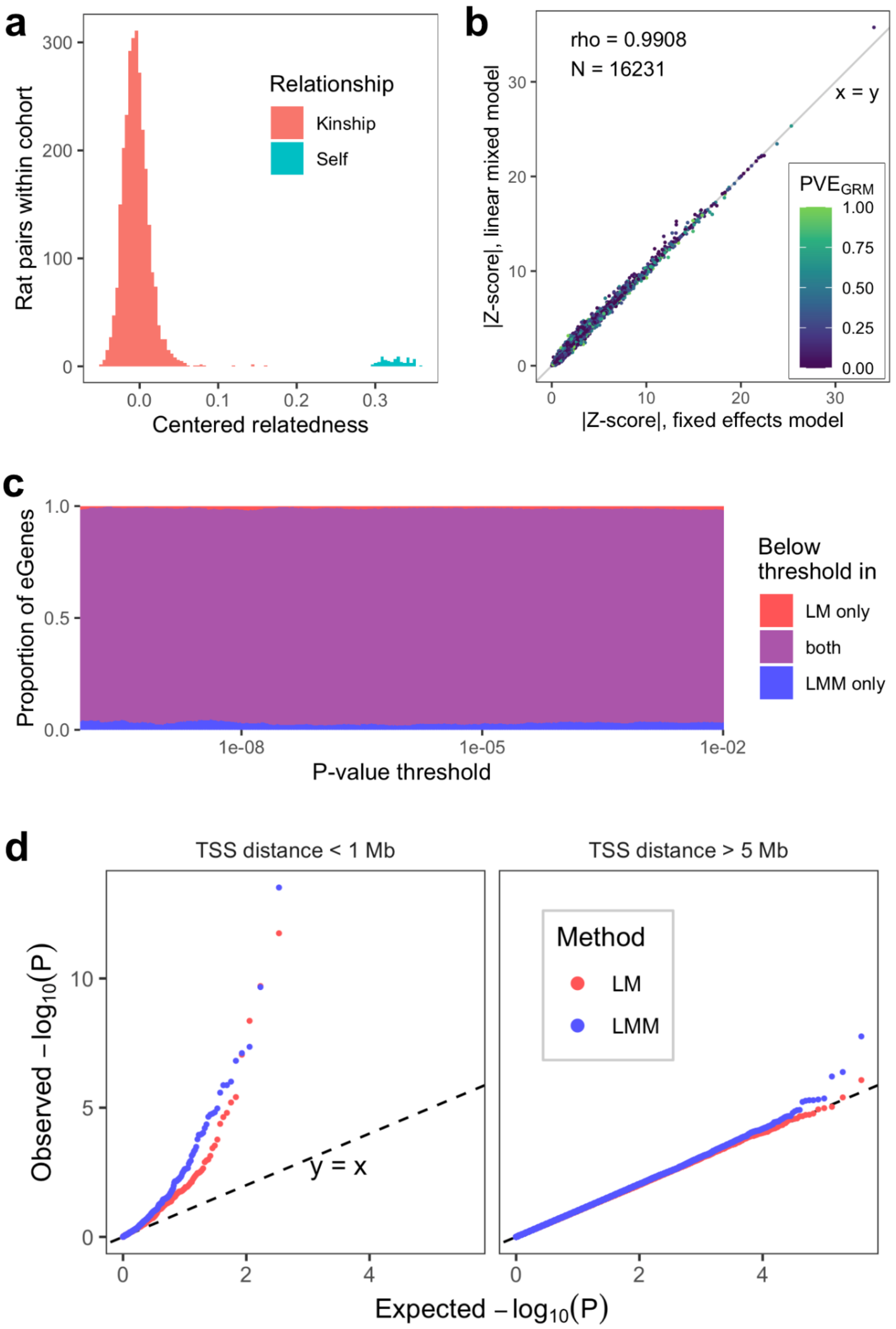


**Impact of random effect term on GEMMA p-values. a** Values from the upper triangle, including the diagonal (Self), of the centered genetic relatedness matrix (GRM) computed from all SNPs in the rats in the nucleus accumbens core dataset. **b** Scatter plots of Z-scores for the top cis-window SNP association per gene for one brain region, nucleus accumbens core. Points are colored by the estimated percent variance explained (PVE) by the GRM. **c** Overlap in the eGene sets below different p-value thresholds. **d** Q-Q plots comparing distributions of -$\log_{10}$(p-values) produced by the LMM and LM methods. A random sample of pruned SNPs were tested per gene per method. Gene-SNP pairs with TSS distance less than 1 Mb are shown separately from those with distances greater than 5 Mb or on different chromosomes, omitting pairs with 1-5 Mb TSS distance.

#### Supplementary Figure 4

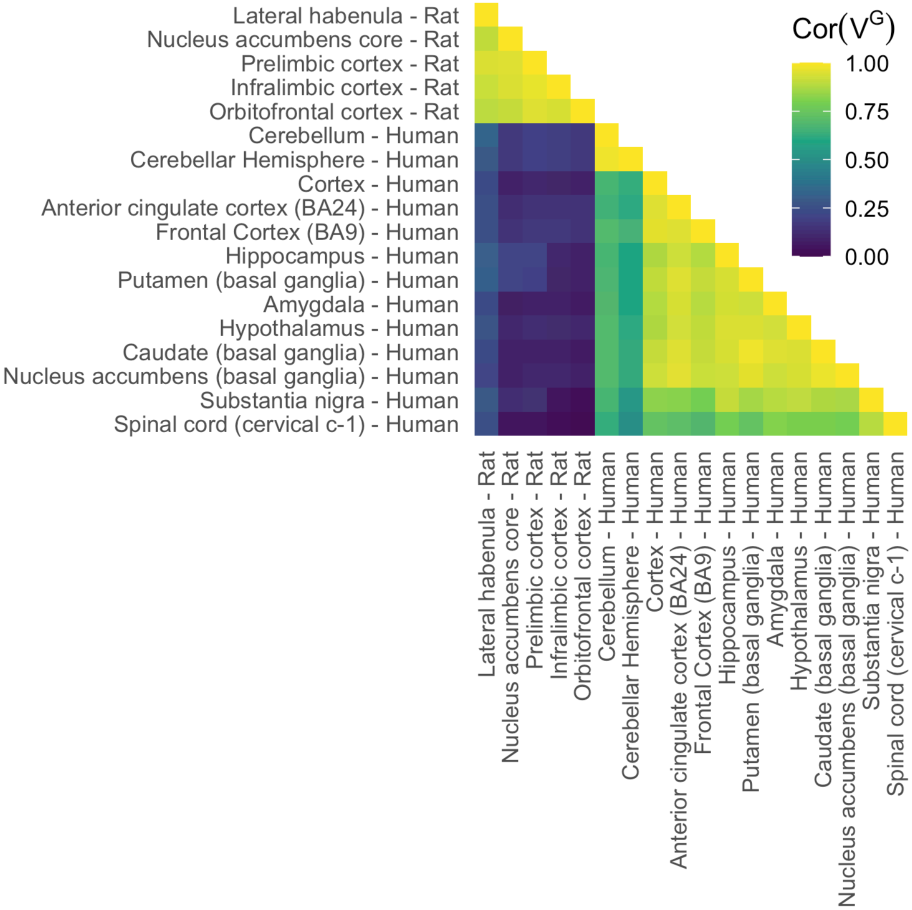


**Heatmap of Pearson correlation of $V^G$ estimates between every pair of human and rat brain tissues.** Ortholog pairs are compared for inter-species tissue pairs, while for tissue pairs from the same species, values for the same gene are compared, including only genes that have an ortholog with a $V^G$ estimate in any tissue.

Supplementary Figure 5

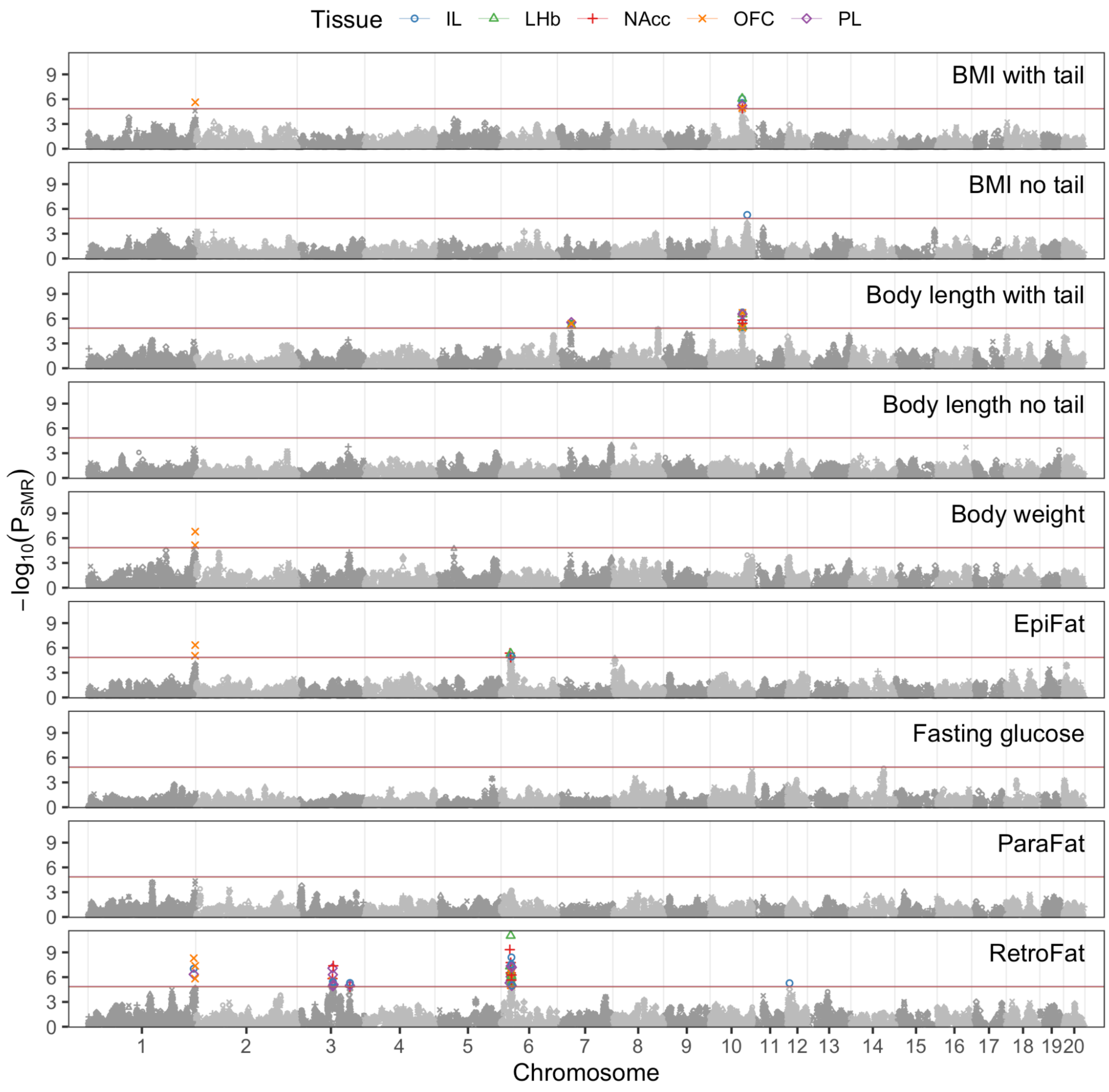


**SMR colocalization P-values for all tissues for each trait.** Each panel overlays p-values for all five eQTL brain tissues tested against the labeled trait. Only significant SNPs are colored by tissue, and the remainder are gray. Bonferroni p-value thresholds for each tissue-trait pair are shown as horizontal lines.
